# Matrigel inhibits elongation and drives endoderm differentiation in aggregates of mouse embryonic stem cells

**DOI:** 10.1101/2023.04.17.537127

**Authors:** Atoosa Amel, Mubeen Goolam

## Abstract

Modelling peri-implantation mammalian development using the self-organising properties of stem cells is a rapidly growing field that has advanced our understanding of cell fate decisions occurring in the early embryo. Matrigel, a basement membrane matrix, is a critical substrate used in various protocols for its efficacy in promoting stem cell growth and self-organization. However, its role in driving stem cell lineage commitment, and whether this effect is driven by biochemical or physical cues is not being clearly defined. Here, we grow embryoid bodies in suspension, Matrigel, and agarose, an inert polysaccharide, to attempt to decouple the physical and biochemical roles of Matrigel and better understand how it drives stem cell differentiation. We show that stem cell aggregates in Matrigel are hindered in their ability to elongate compared to those grown in agarose or in suspension indicating that prohibitive role in self-organisation. Aggregates in Matrigel are also driven to differentiate into endoderm with ectoderm differentiation inhibited. Furthermore, these effects are not due to the physical presence of Matrigel as the same effects are not witnessed in aggregates grown in agarose. Our results thus indicate that Matrigel has a significant and complex effect on the differentiation and morphology of embryoid bodies.

## Introduction

In mammals the establishment of the body plan begins with gastrulation, a process initiated in the posterior region of the embryo through the appearance of the primitive streak (PS) that establishes the formation of the endoderm, mesoderm, and ectoderm [1–4]. Studying these early, key developmental milestones is made challenging in mammals as implantation into the uterine lining renders the embryo inaccessible for direct experimentation and manipulation. However, understanding these early self-organising events remains a fundamental field of study in developmental biology.

Recent advances in stem cell culture techniques have revolutionised the study of early embryonic development as we are now able to study early embryonic milestones using *in vitro* stem cell-based embryo models, so-called ‘stembryos’ [5–13]. A variety of ‘stembryo’ techniques have been developed in a short space of time and have been shown to be able to resemble the blastocyst [14–17], undergo gastrulation [18, 19], develop an anterior-posterior axis [20–22], develop a primitive neural tube [23–25], or even undergo the early stages of somitogenesis [23, 26] and even cardiogenesis [27, 28].

During the development of these protocols one of the critical factors found to drive stem cell self-organisation has been the addition of Matrigel. Matrigel is an extracellular matrix (ECM) derived from Engelbreth-Holm-Swarm mouse sarcoma [29, 30] that contains glycoproteins, proteoglycans, and growth factors and has been found to support embryo culture [31]. Used at different concentration the addition of Matrigel can drive ‘stembryo’ elongation, somitogenesis and neural tube development [23, 26]. These findings suggest that Matrigel plays a key role in driving stem cell differentiation in ‘stembryos’. However, Matrigel has a complex and an unstandardised composition, and its composition can vary from batch to batch [32]. This variability may affect the reproducibility of experiments using Matrigel. Furthermore, Matrigel provides both structural support as well as a host of growth factors and signalling molecules making it unclear whether the advanced morphologies noted in ‘stembryo’ cultures with Matrigel are due to the presence of mechanical or biochemical cues. To uncouple these components, an inert material is needed to evaluate the influence of just mechanical cues on stem cell morphogenesis. Here, we investigate the morphological and gene expression changes induced in embryoid bodies (EBs), the simplest three-dimensional aggregates of stem cells used to study their differentiation, when grown in suspension, Matrigel, and agarose, an inert polysaccharide. In doing so we help to dissect the role of structural support on the fate of mouse embryonic stem (mES) cells.

## Materials and Methods

### mES cell culture

In order to culture 129/Ola mouse embryonic stem (mES) cells, a feeder layer of inactive murine embryonic fibroblast cells was cultured in 6-well plates coated with 0.1% gelatin (Sigma-Aldrich G7041-100G) at a density of 1.6 x 10^4^ cells/cm^2^ in a base medium containing DMEM (1x) +GlutaMAX™ (Gibco 10566016), 15% FBS (Gibco 10493106), 0.05 mM β-mercaptoethanol (Gibco 21985023), and 1% PenStrep (100 U penicillin/0.1 mg/ml streptomycin, Gibco 15140122). Two h before seeding the mES cells, the medium was changed to 2i + LIF medium consisting of base medium supplemented with CHIR99021 (Chiron, 3 µM, Sigma-Aldrich SML1046-5MG), PD0325901 (1 μM, Sigma-Aldrich PZ0162-5MG), and Leukaemia inhibitory factor (10 ng/ml, LIF, Thermo Fisher Scientific A35934). The mES cells were then seeded onto the feeder layer at a density of 7 x 10^3^ cells/cm^2^, and the medium was changed daily, with cells being passaged every other day.

### Cell aggregates in suspension

The aggregate formation method was adopted from Bailie Johnson *et al.* [33]. mES cells colonies were treated with Dispase II (5mg/ml, Sigma-Aldrich D4693-1G) at 37°C for 20 min, followed by inactivation of Dispase and transfer of the cell suspension to a 15 ml tube. The cells were spun down and resuspended in PBS to remove residual medium. Following the PBS washes, the cells were resuspended in N2B27 medium which consists of base medium supplemented with 1% N2 supplement (Gibco 17502048) and 1% B27 plus supplement. (Gibco A3582801). The solution was diluted in N2B27 to give 5×10^4^ cells per 5ml of medium for one 96-well plate. The suspension was then added in 40μl droplets/well to a non-adhesive 96-well U-bottom plate (Greiner Bio-One 650185). The plate was incubated at 37°C and 5% CO_2_ for 48hrs after which a Chiron pulse (CHIR99021, 3µM) was added to the aggregates and the plate was incubated for an additional 24hrs. The following day, the Chiron pulse was removed and fresh N2B27 was added. The medium was changed every day until the required endpoint and not more than 168hrs.

### Cell aggregates on agarose

The 96-well flat bottom plates were coated with 30µl of 1.2% agarose solution (1.2% w/v of Lonza Bioscience 50004 agarose added to deionised water and sent for autoclaving) and left to dry for 10min at RT. mES cell aggregates were made as above, however, instead of adding the cell suspension to 96-well U-bottom plates, 40µl drops were place in each well of the agarose-coated plate. The plate was incubated at 37°C and 5% CO_2_ for 48hrs after which a Chiron pulse (3µM) was added to the aggregates, and they were incubated for an additional 24hrs. The following day, the Chiron pulse was removed and fresh N2B27 was added. The medium was changed every day until the required endpoint and no longer than 168hrs.

### Cell aggregates embedded in Matrigel

mES cells were lifted using Dispase II after a 20 min incubation period and the pellet was washed twice in PBS before counting. The cells were counted, and the appropriate volume was aliquoted and spun down again to give a concentration of 3×10^4^ cells/ml. The pellet was resuspended in 1ml of Matrigel (Corning, 356234). Droplets of 20µl were evenly placed on a 10 cm culture plate and Matrigel was left to solidify for 5 min at 37°C and subsequently covered with N2B27 medium. The Chiron pulse was added to the medium in the dish on day 2 of the culture and removed after 24hrs. The medium was changed every second day until the required endpoint, but not more than 168hrs.

### RNA extraction and cDNA synthesis

RNA was collected from aggregates grown in suspension, agarose or Matrigel 144 hrs post aggregation using the High Pure RNA Isolation Kit (Roche 11828665001) following the manufacturer’s instructions. Due to the limited amount of material obtained from each experiment, two experiments were combined to form one biological replicate. This was done for a total of two biological replicates. The ImProm-II™ Reverse Transcription System (Promega) was utilised for cDNA synthesis. A minimum of 1 µg of RNA was combined with 1 µl of Oligo (dT)15 primer (Promega, C110B) and nuclease-free water to make up a total of 5 µl and incubated at 70°C and 4°C for 5 min each. PCR master mix made up of 6.1 µl nuclease-free water, 4 µl 5x reaction buffer (Promega, M289A), 2.4 µl MgCl2 (Promega, A351H), 1 dNTP mix (Promega, C114B), 0.5 µl Recombinant RNasin® Ribonuclease Inhibitor (Promega, N251A), 1 µl Reverse Transcriptase (Promega, M314A), was added to each sample. Thermal cycling was as follows: 25°C for 5 min, 42°C for 60 min and 70°C for 15 min.

### qPCR

The StepOnePlus™ Real-Time PCR System was used for quantitative PCR with SYBR green PCR Master-Mix (Thermo-Fisher Scientific 4368708). Primer sequences are available in supplementary material Table S1. Briefly, 2 µl of cDNA (diluted 1:1 with nuclease-free water) was mixed with 8 µl master which consisted of: 5 µl SYBR green Master-Mix, 0.4 µl of 10 µM forward and reverse primer mixture (5 µl of 100 µM stock with 40 µl nuclease-free water), and 2.6 µl nuclease-free water. Each reaction had three technical replicates. The run parameters can be found in supplementary material Table S2. *Gapdh* was used as a housekeeping gene to calculate relative expression via the 2-ΔΔCt method. Expression was normalised to cell aggregates cultured in base medium that did not receive a Chiron pulse. Data analysis was performed using MS Excel, while statistical analysis and graph generation were done using GraphPad Prism8.

### Immunofluorescence

Cell aggregates were fixed at 4°C in 4% PFA for 1 h on a shaker. Subsequently, the aggregates were washed three times for 5-10 minutes with PBST (PBS, 0.05% Tween-20) on a rotating shaker and were permeabilised with 0.5% Triton-X-100 in PBS at room temperature for 1 h. After a brief washing step with PBST (Three times for 5-10 minutes), the aggregates were blocked with PBS, 10% FBS, and 0.2% Triton-X-100 at room temperature for 1 h. Primary antibody incubation was performed overnight at 4°C in blocking buffer. The following day, the aggregates were washed three times quickly with PBS, followed by three 20 min washes. Secondary antibody incubation was performed overnight at 4°C in blocking buffer, after which the washing procedure was repeated. Hoechst was added to the 20 min washes following secondary antibody incubation and the samples were left to incubated overnight at 4°C. Prior to imaging, aggregates mounted onto Mowiol drops in glass-bottomed dishes. Antibody details and dilutions used are provided in Table S3.

### Image analysis

The EVOS™ M5000 Imaging System microscope (Thermo Fisher Scientific, USA) was used to capture images at a magnification of 10X. ImageJ was used to optimise the brightness and contrast of all images and for measurements. The outlines of the entire aggregate were traced manually to determine the perimeter, while the minor and major axial lengths were determined using the line tool. The elongation index was calculated by dividing the major axial length by the minor axial length. Multichannel images were taken using a Zeiss LSM 880 Airyscan confocal microscope (Zeiss, Germany) using a 20x or a 40x water-immersion objective. Z-stack slices were spaced at 0.4µm and the final image was deconvolved and displayed at maximum projection using the open-source ImageJ2 platform.

### Statistical analysis

Statistical analyses were caried out using the functions provided by GraphPad Prism 8.4.2 (679). Because of the scarcity of biological material, two biological replicates (each consisting of two pooled experiments) were used for statistical analysis. When appropriate, an unpaired, nonparametric Mann-Whitney test was performed. Alternatively, a Two-way Anova was performed followed by a Fisher’s Least Significant Difference test. In all cases, the error bars represent the mean ± standard error of the mean (mean ± s.e.m.). P values are represented as follows: ns = p≤0.1234, * = p≤0.0332, ** = p≤0.0021, *** = p≤0.0002, **** = p< 0.0001.

## Results

### The effect of Matrigel and agarose on the morphology of mES cell aggregates

To examine the role of physical support in mES cell differentiation, we grew aggregates of ES cells in suspension, on agarose, and in Matrigel (Fig. 1A). Under all conditions embryoid bodies (EBs) were subjected to a pulse of Chiron at 48 h for 24 h and shape changes were quantified over time by measuring the aspect ratio of the aggregates (the longest axis of the aggregate, the major axis, divided by the distance perpendicular to the midpoint of the longest axis, the minor axis) (Fig. 1A). Aggregates were also classified based on visual characteristics and categorised as spherical, ovoid, tear-shaped, budding, or elongating based on their appearance and aspect ratio (Fig. S1). While aggregates grown in suspension (Fig. 1B and E) or agarose (Fig. 1C and F) started to elongate at 72 h, following a pulse of Chiron, those grown in Matrigel primarily retained their spherical or ovoid structures at the same time (Fig. 1D and G). By 144 h, all aggregates in suspension (Fig. 1B and E) and 8/20 of those in agarose (Fig. 1C and F) were able to elongate and resemble the cup-shaped morphology of the embryo. However, aggregates in Matrigel retained a more oval shape by the end of the culture period and did not elongate as easily (Fig. 1D and G). It was also noted that agarose aggregates had a unique, temporary balloon-like morphology (Fig. S1) that appeared at 24 h (16/20) and were mostly no longer present by 96 h (1/20) (Fig. 1F).

**Fig. 1:**
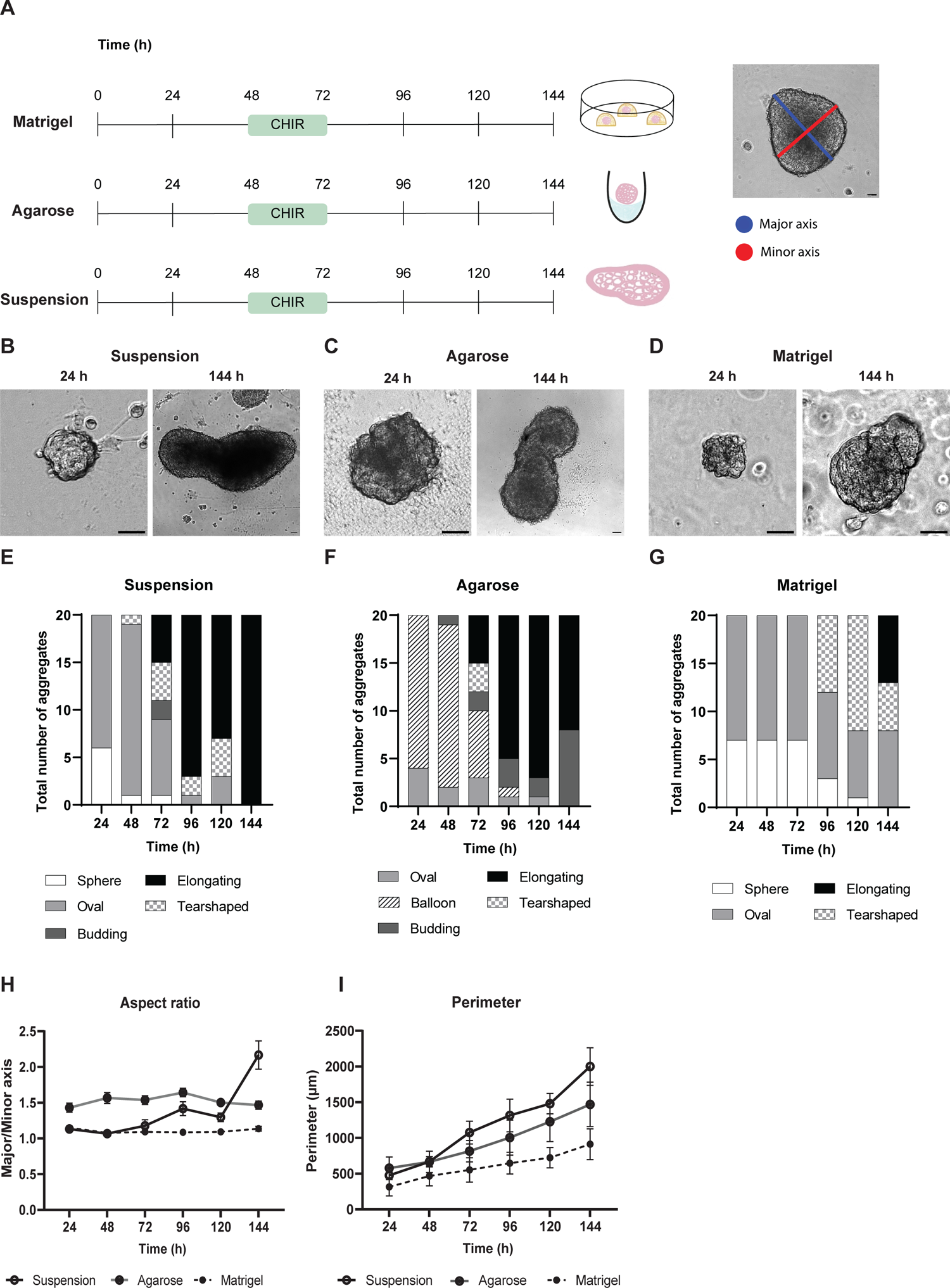
Comparative analysis of the effect of various physical constraints on ES cell aggregate morphology. (A) ES cell aggregates were cultured in suspension, on agarose or embedded in Matrigel. All aggregates received a 24 h Chiron pulse on the third day of culture. To calculate the elongation of aggregates, the major axis (the longest axis; blue line) was measured and divided by the minor axis-the line perpendicular to the major axis at its midpoint (red line). (B) Morphology of aggregates in suspension. Aggregates had elongated by 144 h. (C) Morphology of aggregates cultured on agarose. At 24 h the aggregates had already showed signs of asymmetry and by 144 h the aggregates had elongated. (D) Morphology of aggregates embedded in Matrigel. The aggregates were relatively smaller than the other two conditions and did not elongate significantly. (E) Characteristics of n = 20 aggregates grown in suspension for the period of 144 h. Major morphological changes occurred at 72 h and at 144 h all the aggregates had elongated. (F) Characteristics of n = 20 aggregates grown on agarose for the period of 144 h. The aggregates had a unique balloon-like morphology in the first 96 h of culture and by 144 h some aggregates were still in the budding stage and had not elongated fully. (G) Characteristics of n = 20 aggregates grown in Matrigel for the period of 144 h. Aggregates had a delayed growth and only showed signs of asymmetry at 96 h. (H) Aspect ratio of aggregates in suspension (black line, empty circle), on agarose (grey line, dotted circle) and in Matrigel (dotted line, solid circle) for n = 20 aggregates per condition. (I) Perimeter of aggregates in suspension (black line, empty circle), on agarose (grey line, dotted circle) and in Matrigel (dotted line, solid circle) for n = 20 aggregates per condition. Error bars indicate mean ± SEM. The aspect ratio within the same condition between two consecutive time points was evaluated for significance. An unpaired, nonparametric Mann-Whitney test was performed and * = p≤0.0332, ** = p≤0.0021, *** = p≤0.0002, **** = p< 0.0001. Scale bar: 100 µm.

Aspect ratio analyses revealed that suspension aggregates were the only group that underwent a rapid period of elongation at 72 h and an additional increase in axial length at 144 h (Fig. 1H). Interestingly, there was a dip in aspect ratio at 120 h under both suspension and agarose conditions, meaning that the aggregates grew in width rather than length. The suspension aggregates had the highest rate of growth in perimeter, followed by the agarose aggregates, and the Matrigel embedded aggregates showed the smallest increase in perimeter during the 144 h period (Fig. 1I).

### Matrigel drives endoderm differentiation in mES cell aggregates

Physical support from the maternal environment plays an essential role in early embryonic development during induction of the PS, when cells first undergo an epithelial-to-mesenchymal transition (EMT) and form mesodermal and endodermal progenitors [34]. In this study, we sought to determine the effects of physical constraints on the development of the PS and in driving EMT in EBs. To investigate this, we measured the expression of EMT and PS markers using qPCR in EBs grown in suspension, agarose, and Matrigel with a Chiron pulse, and compared this to EBs grown in suspension without any exposure to a Wnt agonist.

The posterior epiblast marker *Wnt3* was significantly increased in suspension aggregates compared to agarose and Matrigel aggregates while there was little variation in the expression of *ß-catenin* (Fig. 2). *Brachyury,* a marker of the PS and mesoderm, was also upregulated at 144 h under suspension conditions, with no increase in expression noted in agarose or Matrigel conditions (Fig. 2). However, another PS and mesoderm marker, *Nodal*, was significantly up-regulated in both agarose and Matrigel conditions (Fig. 2). The mesendoderm marker *Eomes* was up-regulated in EBs in suspension, as well as in Matrigel (Fig. 2), while another mesendoderm marker *Mixl1* was significantly up-regulated significantly in suspension and agarose conditions but not in Matrigel (Fig. 2). Interestingly, the early endoderm marker, *Gata6*, had increased levels of expression in Matrigel aggregates compared to the control (although not significantly), while a marker of the definitive endoderm marker, *Sox17,* was significantly increased in Matrigel compared to the other conditions, suggesting that Matrigel may be directing differentiation towards the endoderm lineage. A key EMT marker, *Snai1* was down-regulated significantly under suspension and agarose conditions compared to the control (Fig. 2) and *E-cadherin*, an epithelial marker, was down-regulated in suspension compared to the other two conditions (Fig. 2). The anterior epiblast marker *Pou3f1* had higher levels of expression in all three conditions compared to controls, albeit with a significant increase only observed in suspension (Fig. 2). Similarly, *Slc7a3*, another anterior epiblast marker, had increased expression levels in all three conditions compared to controls with a significant increase in suspension and agarose conditions (Fig. 2). The anterior/neuroectoderm marker *Sox2* had an increased relative level of expression in the three conditions, while *Pax6*, a late neurectoderm marker, was only up-regulated in suspension aggregates. This suggests that neuroectoderm formation may be at a more advanced stage in suspension aggregates exposed to a Chiron pulse.

**Fig. 2:**
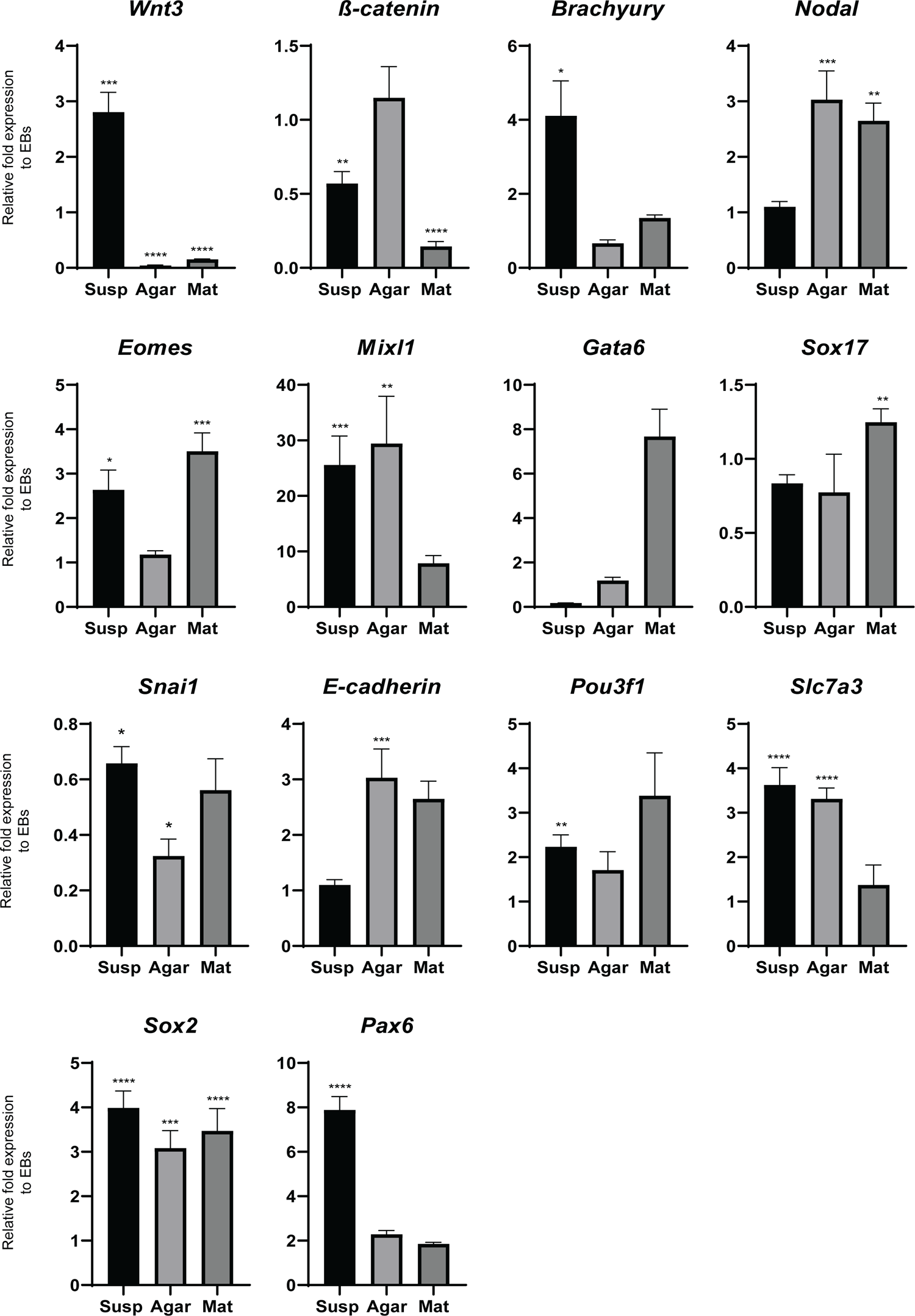
Effects of various physical constraints on gene expression of aggregates. QPCR was performed on reverse transcribed RNA extracted at 144 h. Primitive streak markers: *Wnt3, ß-catenin, Brachyury* and *Nodal.* Mesendoderm markers: *Eomes* and *Mixl1*. Endoderm markers: *Gata6* and *Sox17*. EMT markers: *Snai1* and *E-cadherin*. Anterior/neurectoderm markers: *Pou3f1, Slc7a3, Sox2* and *Pax 6.* Relative fold expression to cell aggregates without a Chiron pulse. Error bars indicate mean ± SEM. An unpaired, nonparametric Mann-Whitney test was performed and * = p≤0.0332, ** = p≤0.0021, *** = p≤0.0002, **** = p< 0.0001.

### Aggregates in Matrigel form pro-amniotic-like cavities

The effects that physical constraints had on mesoderm and endoderm gene expression next led us to investigate the spatial expression of the mesoderm marker Brachyury and the endoderm marker Sox17 in our aggregates using immunofluorescence. When in suspension very few aggregates expressed Brachyury. When they did, this expression was either restricted to one pole of the aggregate (Fig. 3A, 1/50) or dispersed throughout the entire aggregate (Fig. 3B, 2/50). A highly similar pattern was observed in the agarose aggregates with only a few aggregates being positive for Brachyury, and when they were, this staining was either localised (Fig. 3C, 2/46) or dispersed (Fig. 3D, 2/46). Aggregates in Matrigel, however, have significantly more Brachyury positive aggregates with 18/60 showing a dispersed pattern of expression Fig. 3E). Interestingly, all of these also developed a Brachyury lined lumen in their centers (Fig. 3F, 18/60). All aggregates in Matrigel were found to have formed a similar cavity, even those without any Brachyury-positive staining. The cells surrounding these cavities appeared as epithelial-like and were brightly stained with E-cadherin (Fig. 3G). The suspension and agarose aggregates clearly lacked such cavities, but they had regions that were negative for E-cadherin (Fig. 3H and 3I) suggesting that EMT has occurred.

**Fig. 3:**
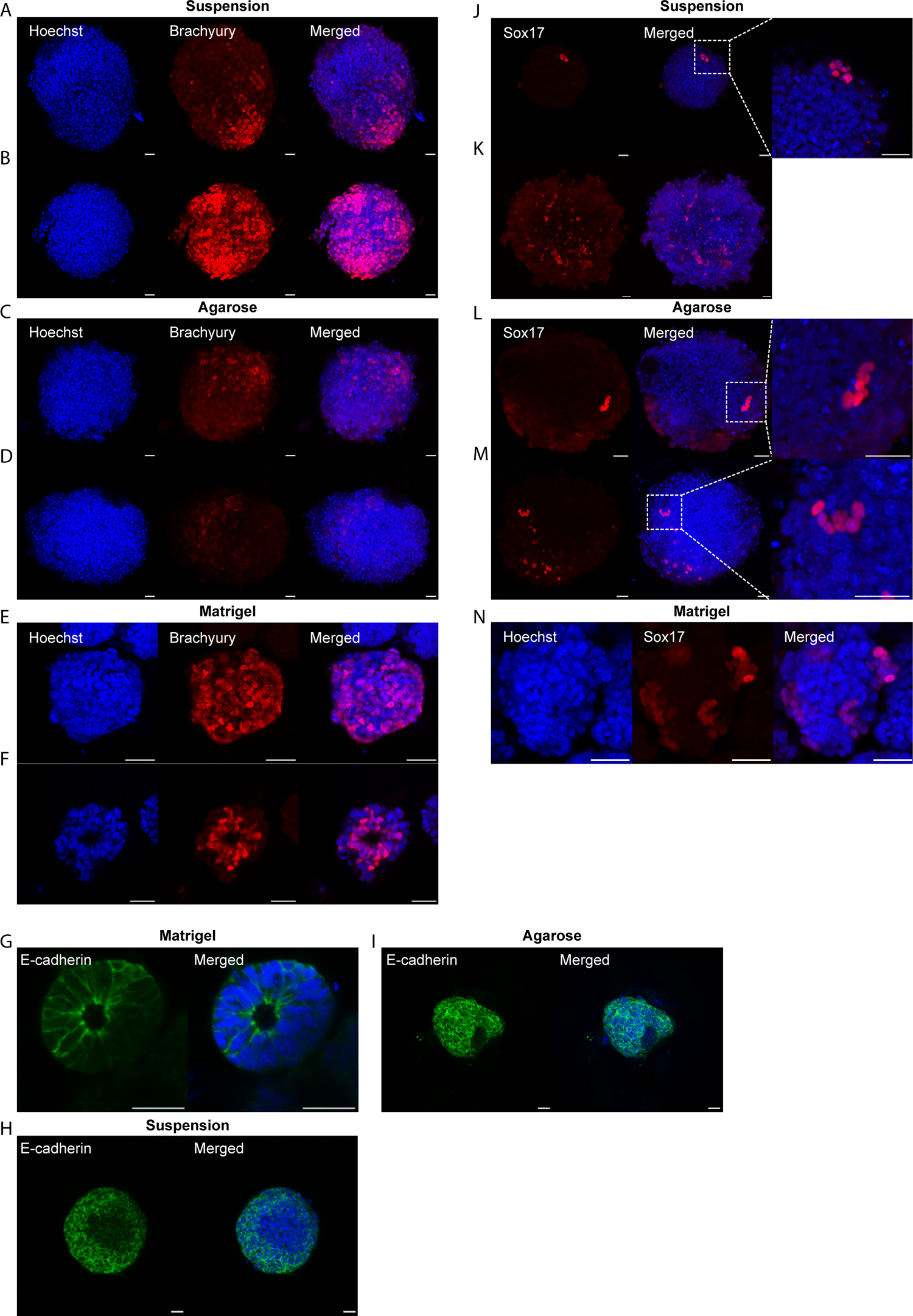
Immunofluorescence staining of Brachyury, Sox 17 and E-cadherin in aggregates cultured under various physical constraints. (A) Localised Brachyury positive cells at the elongating tip of suspension aggregates. (B) Brachyury positive cells spread throughout suspension aggregates that remained spherical. (C) Localised Brachyury positive cells in agarose aggregates. (D) Dispersed Brachyury-positive cells in agarose aggregates. (E) Brachyury positive cells dispersed throughout Matrigel aggregates. (F) Brachyury positive cells lining the lumen of Matrigel aggregates. (G) Epithelial-like cells surrounding the cavity in Matrigel aggregates stained positive for epithelial marker E-cadherin. (H) Suspension aggregates stained for E-cadherin had regions with reduced expression that may represent EMT. (I) Agarose aggregates stained for E-cadherin had regions with reduced expression that may represent EMT. (J) Pockets of Sox17 positive cells in the periphery of suspension aggregates. (K) Sox17 positive cells dispersed across the surface of suspension aggregates. (L) Pockets of Sox17 positive cells in aggregates cultured on agarose. (M) Sox17 positive cells dispersed across the surface of aggregates grown on agarose. (N) Sox17 positive cells in dispersed groups across the surface of aggregates in Matrigel. Scale bar: 25µm.

Endoderm staining was observed infrequently in all conditions investigated. In suspension aggregates, it was noted as either small pockets of Sox17-positive cells near the periphery (Fig. 3J, 7/46) or dispersed across the surface of the aggregate (Fig. 3K, 7/46). A similar expression pattern was observed in agarose aggregates with small pockets of Sox17-positive cells (Fig. 3L, 3/49) or a dispersed expression pattern (Fig. 3M, 5/49). Aggregates in Matrigel were noted to express Sox17 in dispersed groups across the surface of the aggregate (Fig. 3N, 5/60).

## Discussion

The emergence and subsequent development of ‘stembryo’ culture systems has generated a non-invasive, scalable, and novel way to investigate early mammalian embryonic morphogenesis and cell fate decisions gastrulation and germ layer formation in mammals. One of the key aspects missing from ‘stembryo’ structures is the presence of physical support provided by the ECM normally generated by the basement membrane of the visceral endoderm. To address this issue, many of these systems incorporate the commercial basement membrane substitute Matrigel. Although Matrigel can promote stem cell self-organisation and provides some degree of mechanical constraint, it is ill-defined and suffers from lot-to-lot variability. In addition, its effect on organoid structure and differentiation is influenced by both mechanical and biochemical cues, and the specific component/s responsible for the observed morphologies remain unknown. Matrigel also does not support any controlled modifications of its stiffness or components, and thus there is a need to use an inert and manipulatable scaffold to separate the effects of physical and chemical cues on stem cell self-organisation in this burgeoning field. In this study, we used agarose, an inert polysaccharide, to test the role of mechanical cues on the fate of embryoid bodies and to try to create similar physical constrains to those provided by the uterus.

We found that EB morphology was significantly influenced by the physical constraints present during culture, with suspended aggregates exhibiting more effective growth and elongation than counterparts grown in the presence of Matrigel or agarose. At 48 h, some suspension and agarose aggregates began to exhibit tear-shaped or budding structures, indicating an intrinsic breaking of symmetry. At 72 h, there was a sudden increase in the variety of morphologies, including balloon-like structures, which were observed as intermediates in the elongation process. Interestingly, we found that the presence of Matrigel inhibited elongation, causing aggregates to maintain their oval or spherical shape for longer. When we measured aspect ratios, we found that aggregates in suspension underwent a drop in aspect ratio at 120 h, followed by a sudden increase at 144 h. This suggests that cell proliferation occurs before elongation, along the major axis. Aggregates grown in agarose exhibited a similar dip and rise in aspect ratio, but the same pattern was not found in aggregates grown on Matrigel. Overall, we found that early self-patterning events in stem cell aggregates in culture are hindered by the presence of Matrigel. These aggregates neither grow nor elongate as effectively as those in suspension or in agarose, with Matrigel playing a prohibitive role in self-organisation.

Previous studies have shown that after 144 h in culture ‘stembryo’ should start to undergo a gastrulation-like process [20, 23, 35, 36]. Although we noted elongation in EBs in both suspension and agarose conditions by this time point, there were marked differences in the expression of key markers between the two conditions. Aggregates in suspension had increased levels of *Brachyury* suggesting that the PS had been initiated. In contrast, aggregates grown in agarose did not have this same up-regulation in *Brachyury* expression. They did, however, have up-regulated *ß-catenin,* an upstream regulator of *Brachyury*. This could indicate a delay in the gastrulation process due to the differences in the physical properties of the environments as the more restrictive agarose may present a less conducive environment for elongation and subsequent gastrulation. Aggregates in Matrigel had similarly low levels of *Brachyury* and *ß-catenin,* further supporting the theory that physical support hinders stem cell driven *in vitro* gastrulation-like events. Aggregates in suspension also had the lowest relative levels of *E-cadherin*, an epithelial marker, and the highest levels of mesenchymal marker *Snai1,* suggesting that these aggregates are likely undergoing EMT, a critical process in gastrulation. This is further support for suspension aggregates being more developmentally advanced than aggregates in either agarose or Matrigel.

However, the expression patterns of *Nodal* and *Wnt3* in our aggregates indicate that the developmental landscape in our conditions is more complex than this. *Nodal* and *Wnt3* are known to work together to induce the expression of *Brachyury* and *Eomes*, another marker of gastrulation [1]. We observed that *Nodal* and *Wnt3* have opposing expression patterns in suspension aggregates, with elevated levels of *Wnt3* and low levels of *Nodal*. Interestingly, the opposite pattern, high levels of *Nodal* and relatively low levels of *Wnt3* was observed in aggregates in agarose and Matrigel. These results would indicate that the aggregates under these conditions are more advanced in terms of mesoderm differentiation. As *Eomes* has been shown to precede *Brachyury* expression [37], and based on the high levels of *Eomes* we can assume that *Brachyury* expression would increase in both suspension and Matrigel aggregates if cultured further. As both aggregates in Matrigel and agarose show a similar pattern of expression in this case, we hypothesise that it is principally mechanical cues driving this expression pattern and this warrants further investigation.

Analysing the expression of key markers of PS formation and EMT transition, it appears that our aggregates most closely resemble mouse embryos at E5.5-E6.5, between the emergence of the germ layers and the formation of the anterior-posterior axis (A-P axis), which occurs at approximately 120-144 h of development[6]. It is noted that this is significantly delayed compared to other ‘stembyro’ culture processes, where aggregates can undergo a gastrulation-like process after 96 h and exhibit similarities to the E8.5 mouse embryo [20, 23, 35, 38].

Despite this apparent developmental delay, suspension aggregates had high levels of anterior/neuroectoderm markers, *Pou3f1, Slc7a3, Sox2*, indicating the initiation of the anterior-posterior(A-P) axis. We observed a distinct increase in in *Pax6,* a late neuroectoderm marker, in suspension aggregates similar to what is observed in E8.5 mouse embryos and gastruloids (one of the most robust ‘stembyro’ systems) at 120 h [38]. This is possibly due to the high levels of *Sox2* present in mES cells prior to differentiation, allowing rapid neural differentiation [39], provided they were able to differentiate without any confounding effects, such as physical constraints.

Aggregates in Matrigel demonstrated the highest expression of endoderm markers compared to the other two conditions. *Gata6* is typically a marker for cardiac mesoderm and DE progenitors [40], while *Sox17* is a well-known DE marker [41]. *Gata6* mRNA was up-regulated in Matrigel aggregates, followed by up-regulation of *Sox17*, indicating that this condition likely favoured endoderm formation. It is unclear if this preference is due to physical restrictions imposed by Matrigel or the presence of certain signalling factors. However, evidence suggests that laminin, a major component of Matrigel, can direct human ES cells toward the endoderm lineage [42]. Furthermore, studies have demonstrated that culturing EPI stem cells in Matrigel results in up-regulated levels of endoderm-related transcription factors such as Sox17 [43], further supporting the notion that Matrigel influences lineage commitment towards endoderm.

Immunofluorescent staining for Brachyury revealed that the positive cells were located exclusively at the elongated pole of the aggregates in both suspension and agarose, as observed in other studies [20, 35, 38, 44], although the frequency of this pattern was significantly lower than reported by others. Aggregates in Matrigel, however, had many more aggregates that were Brachyury-positive, and these cells were principally in the centre of the aggregate surrounding the lumen. Studies have demonstrated that Matrigel can increase the expression of Wnt3 antagonists such as Dkk1 and Sfpr1 [45]. Therefore, it is plausible that cells located in the periphery of our Matrigel aggregates express Wnt3 agonists, resulting in the localisation of Wnt3 activity in the centre of the aggregates, which leads to the up-regulation of Brachyury expression. By embedding aggregates in Matrigel, we were able to observe cavity formation that resembled the pro-amniotic cavity. This was likely due to the cells pulling apart, facilitated by E-cadherin localisation around the membrane. Matrigel is used as a substitute for ECM derived from extra-embryonic tissue [18] and the interaction between cells and Matrigel via integrin receptors facilitates cell polarisation and cavity formation [46]. In the embryo, cavity formation is mediated by the interaction between laminin and the ß1-integrin receptor between embryonic and extra-embryonic tissue [46]. The same mechanism may operate in Matrigel aggregates, allowing them to form pro-amniotic cavity-like structures. To test the establishment of polarity in our aggregates, future work should involve immunofluorescence staining for aPKC and Par6 [47, 48].

Overall, our results suggest that Matrigel has a significant and complex effect on the differentiation and morphology of mES cells in culture. It can hinder stem cell self-organisation and neural differentiation, drive endoderm differentiation, and induce the formation of a pro-amniotic cavity. Its effects are not driven simply by the mechanical force it provides as its effects on stem cells are not mimicked by using agarose. Although Matrigel is a critical component of ‘stembryo’ cultures, caution should be taken when viewing it as a substitute for an ECM. Future research in the field should focus on finding a more defined and manipulatable replacement for Matrigel in order to advance the field and the accuracy of *in vitro* embryo models.

## Author contributions

M.G. conceived the project. A.A carried out the experiments. A.A. and M.G. wrote the paper.

## Declaration of competing interest

The authors have no financial or competing interests to declare.

## Data availability

Data will be made available upon request.

## Acknowledgements

We thank Frank Brombacher for the mES cells and Thulisa Mkatazo for assistance with generating inactivated murine embryonic fibroblasts.

## Funding

This work was supported by the South African National Research Foundation (M.G., Competitive Support for Unrated Researchers - 129399, A.A. Postgraduate MSc Scholarship) and the University of Cape Town (M.G., University Research Committee Start Up Grant, Faculty of Health Science Start Up Emerging Researchers Award, Research Development Grant, Enabling Grant Seeker Excellence Award).

## Notes

### Competing Interest Statement

The authors have declared no competing interest.

