## Supplementary Material for "Matrigel inhibits elongation and drives endoderm differentiation in aggregates of mouse embryonic stem cells"

**Table S1: Primers**

| Gene | Forward sequence (5' to 3') | Reverse sequence (5' to 3') | Amplicon size (bp) |
|---|---|---|---|
| *Gapdh* | AGGTCGGTGTGAACGGATTTG | TGTAGACCATGTAGTTGAGGTCA | 123 |
| *Sox2* | GCGGAGTGGAAACTTTTGTCC | CGGGAAGCGTGTACTTATCCTT | 157 |
| *Sox17* | GATGCGGGATACGCCAGTG | CCACCACCTCGCCTTTCAC | 136 |
| *Gata6* | GTGGTCGCTTGTGTAGAAGGA | TTGCTCCGGTAACAGCAGTG | 105 |
| *Brachyury* | GCTGGATTACATGGTCCCAAG | GGCACTTCAGAAATCGGAGGG | 158 |
| *Mixl1* | GTCTTCCGACAGACCATGTACC | CCCGCCTTGAGGATAAGGG | 160 |
| *Pou3f1* | TTCAAGCAACGACGCATCAA | TGCGAGAACACGTTACCGTAGA | 86 |
| *Slc7a3* | TTCTGGCCGAGTTGTCTATGTTTG | AGTGCGGTTCTGTGGCTGTCTC | 190 |
| *E-cadherin* | CAGGTCTCCTCATGGCTTTGC | CTTCCGAAAAGAAGGCTGTCC | 175 |
| *Nodal* | CCTGGAGCGCATTTGGATG | ACTTTTCTGCTCGACTGGACA | 155 |
| *Snai1* | CTTGTGTCTGCACGACCTGT | ACATCCGAGTGGGTTTGGAG | 167 |
| *Pax6* | TACCAGTGTCTACCAGCCAAT | TGCACGAGTATGAGGAGGTCT | 194 |
| *ß-catenin* | ATGGAGCCGGACAGAAAAGC | CTTGCCACTCAGGGAAGGA | 108 |
| *Eomes* | TCGCTGTGACGGCCTACCAA | AGGGGAATCCGTGGGAGATGGA | 210 |
| *Wnt3* | CTCGCTGGCTACCCAATTTG | CTTGACACCTTCTGCTACGCT | 165 |

**Table S2: StepOnePlus™ Real-Time PCR run parameters for a 2hr run.**

| Stage | Step | Temperature (°C) | Duration |
|---|---|---|---|
| Holding | 1 | 95 | 10 min |
| Cycling (40 cycles) | 1 | 95 | 15 s |
| | 2 | 60 | 1 min |
| Melt Curve (continuous) | 1 | 95 | 15 s |
| | 2 | 60 | 1 min |
| | 3 | 95 (with a ramp rate of 2.8%) | 15 s |

**Table S3: Antibodies**

| Antibody (Source) | Supplier | Cat. # | Dilution |
|---|---|---|---|
| Hoechst | Thermo Fisher Scientific | H1399 | 1:1000 |
| Anti-Human/mouse Brachyury (Goat) | R&D systems | AF2085 | 1:100 |
| Anti-Mouse E-cadherin (Mouse) | Takara Bio Inc. | M108 | 1:500 |
| Anti-Human SOX17 (Goat) | R&D systems | AF1924 | 1:100 |
| Cy3 (Donkey anti-goat) | AEC-Amersham SOC Ltd | 705 166 147 | 1:500 |
| Alexa 488 (Donkey anti-mouse) | AEC-Amersham SOC Ltd | 715 546 150 | 1:500 |

A

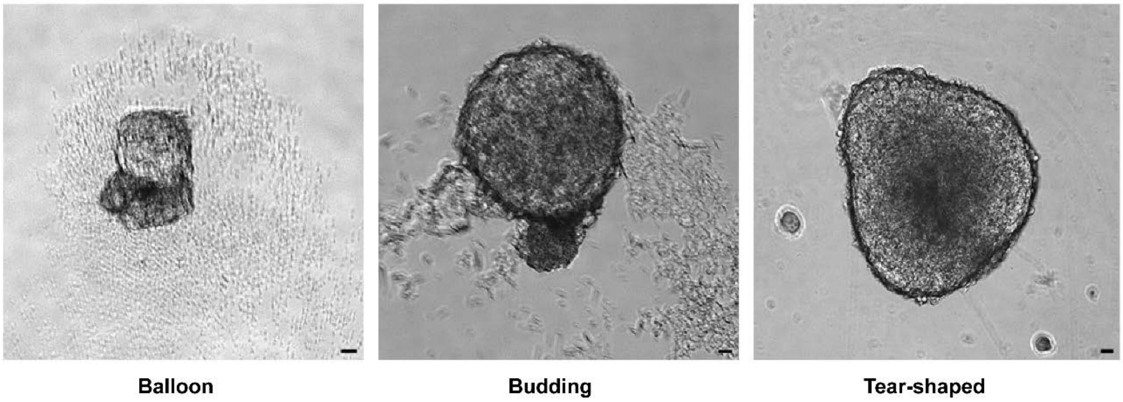


B

| Shape | Aspect ratio range |
|---|---|
| Spherical | 1.0 < X < 1.01 |
| Oval/Budding | 1.01 < X <1.35 |
| Tear-shaped | 1.15 < X < 1.4 |
| Elongating | X > 1.3 |

**Fig. S1: Physical characteristics of aggregates grown under various physical constraints.** (A) Balloon, budding and tear-shaped morphologies observed in aggregates. The Balloon morphology was unique to aggregates cultured on agarose. (B) Aspect ratio used to classify aggregates. Aggregates were categorised as spherical, ovoid, tear-shaped, budding, or elongating based on their appearance and aspect ratio.
